# Cytoplasmic Fluidity and the Cold Life: Proteome Stability is Decoupled from Viability in Psychrophiles

**DOI:** 10.1101/2025.06.24.661299

**Authors:** Beatrice Caviglia, Stepan Timr, Marianne Guiral, Marie-Thérèse Giudici-Orticoni, Tilo Seydel, Christian Beck, Judith Peters, Fabio Sterpone, Alessandro Paciaroni

**Affiliations:** Department of Physics and Geology, University of Perugia, via Alessandro Pascoli, 06123 Perugia, Italy; Laboratoire de Biochimie Théorique (UPR9080), CNRS, Université de Paris Cité, 75005 Paris,France; Institut de Biologie Physico-Chimique, Fondation Edmond de Rothschild, 13 Rue Pierre et Marie Curie, 75005 Paris, France; J. Heyrovský Institute of Physical Chemistry, Czech Academy of Sciences,182 00 Prague, Czechia; Laboratoire de Bioénergétique et Ingénierie des Protéines, BIP, CNRS, Aix-Marseille Université, 13400 Marseille, France; Institut Laue-Langevin, 71 avenue des Martyrs CS 20156, 38042 Grenoble, France; Université Grenoble Alpes, CNRS, Laboratoire Interdisciplinaire de Physique, 140 Rue de la Physique, 38402 Saint-Martin-d’Hères, France; Institut Universitaire de France, 103 Saint-Michel, 75005 Paris, France

## Abstract

Protein diffusion, critical for cellular metabolism, occurs in the highly crowded cytoplasm. Understanding how this dynamics changes when organisms are adapted to different thermal niches is a fundamental challenge in microbiology and biophysics. In *Escherichia coli*, protein diffusion undergoes a pronounced slowdown at temperatures near cellular death, coinciding with the early stages of unfolding. To determine whether this phenomenon is universal, we investigated psychrophilic and hyperthermophilic bacteria. In both species, a marked diffusion slowdown takes place at the onset of proteome melting. However, while the dynamic arrest is associated with the thermal death point for the hyperthermophilic proteome, the psychrophilic proteome maintains substantial mobility well beyond the cellular inactivation. The decoupling between metabolic viability and proteome dynamics and stability suggests that the functional processes of psychrophilic bacteria are extremely temperature sensitive. This finding echoes the behaviour of psychrophilic enzymes, manifesting a large temperature gap between optimal activity and unfolding. Protein diffusion is optimised to maintain functional fluidity at the organism’s working conditions, but its temperature dependence is controlled by the proteome folded state. Our findings redefine the relationship between cytoplasmic dynamics, proteome stability, and bacterial survival in cold environments.

## I. INTRODUCTION

Biological life has evolved in response to environmental conditions, which have both shaped and constrained its development. Among these factors, temperature plays a pivotal role in regulating the growth of diverse organisms on Earth. A wide range of extremophiles - organisms living under extreme conditions - were discovered and studied in the last century [1]. *Planococcus halocryophilus* and *Methanopyrus kandleri*, for example, represent the most extreme cases of temperature adaptation known to date, with the former growing at −15°C and remaining metabolically active at 25°C [2], while the latter thriving at a record-high temperature of −122°C [3]. These organisms have drawn significant interest for their potential applications in diverse fields, including biotechnology and cancer research. The widely used biotechnological technique Polymerase Chain Reaction (PCR), for example, relies on thermostable enzymes derived from thermophilic and hyperthermophilic bacteria for DNA amplification [4, 5]. Thermostability also plays a crucial role in cancer treatment, where hyperthermia is used to selectively kill cancerous cells through exposure to elevated temperatures [6]. Additionally, current research is exploring the potential for life on extraterrestrial habitats, such as Mars, Venus, and icy moons, where extreme conditions prevail, making extremophiles key models for astrobiology and the search for life beyond our planet [7, 8]. Finally, climate change drives the selection of bacterial species that are more resistant to high temperatures, which correlates with a higher propensity for antibiotic resistance [9].

Defining the limits of life’s growth is essential not only for practical applications but also for advancing our understanding of evolutionary biology. However, the fundamental mechanisms underlying cellular thermostability remain largely elusive. Deciphering the factors that drive cell death at high temperatures and identifying what enables certain bacteria to endure extreme heat more effectively than others poses major scientific challenges.

Temperature impacts all bio-molecular components of the cell, as well as biophysical processes within a cell. High temperatures affect the structures of proteins, DNA, RNA, and cell membranes [10], making it difficult to unravel the molecular events that ultimately lead to cell death [11–14]. Among macromolecules, proteins are the most abundant yet least stable biomolecules in the cell, making their thermal sensitivity a key factor in regulating temperature-dependent cellular activities [15].

Recently, we showed that the unfolding of just a small fraction of the *E. coli* proteome is able to give rise to a dynamic arrest around the thermal cell death [16]. This phenomenon is of key biological relevance, as diffusive dynamics plays a fundamental role in cell growth by ensuring the proper spatial and temporal organization of cytoplasmic components and regulating solute partitioning during cell division [17]. Moreover, it governs the mobility of intracellular molecules, setting the constraints for essential molecular interactions and biological reactions [18, 19].

Therefore, a fundamental question arises: is the arrest of protein dynamics at cell death a universal phenomenon, or is it exclusive to mesophiles like *E. coli* ? If psychrophiles and (hyper)thermophiles exhibit similar or distinct diffusive slowdowns at their respective cell death temperatures, it could unveil key principles underlying cellular thermostability and evolutionary adaptation. To address this question, we combined molecular dynamics (MD) simulations and quasi-elastic neutron scattering (QENS) experiments to systematically compare the proteome dynamics of three bacterial species adapted to different thermal environments: the psychrophile *P. arcticus* (PA), the mesophile *E. coli* (EC), and the hyperthermophile *A. aeolicus* (AA).

For *A. aeolicus*, we observe a dynamic arrest closely mirroring the behaviour of *E. coli*, occurring near its corresponding cell death temperature of ~ 370 K. In contrast, *P. arcticus* exhibits a markedly different behavior, with protein diffusion abruptly declining well above its lethal threshold of ~ 295 K. In all three organisms, the arrest of protein diffusive dynamics is driven by the unfolding of a small fraction of the proteome. However, in *P. arcticus*, this arrest is decoupled from cell death, which instead likely results from the loss of biological activity in key enzymes already within the low-temperature range.

## II. RESULTS

### A. Global Diffusion of Bacterial Proteins Slows Down at Critical, Species-Dependent Temperatures

We utilized quasi-elastic neutron scattering (QENS) experiments to investigate the diffusive dynamics of an average bacterial protein on the nanosecond timescale. The underlying hypothesis is that the observed scattering primarily originates from the incoherent contribution of hydrogen atoms within bacterial proteins [20], which constitute the predominant biomolecular species in the bacterial cytoplasm [21, 22] (see Methods section IV B).

QENS experiments were performed on the IN16B spectrometer at the Institute Laue Langevin [23] on PA, EC, and AA bacteria, across an extended temperature range to include the respective thriving conditions and the upper thermal stability limit.

The incoherent dynamic structure factor, *S*(*Q, E*), obtained from the experiments and shown in Fig. 1a, can be described as a combination of two distinct contributions: local dynamics, which include vibrations and local diffusive motions within the interior of proteins, and global dynamics, which encompass the rotational and translational diffusive motions of proteins as a whole [24].

**FIG. 1:**
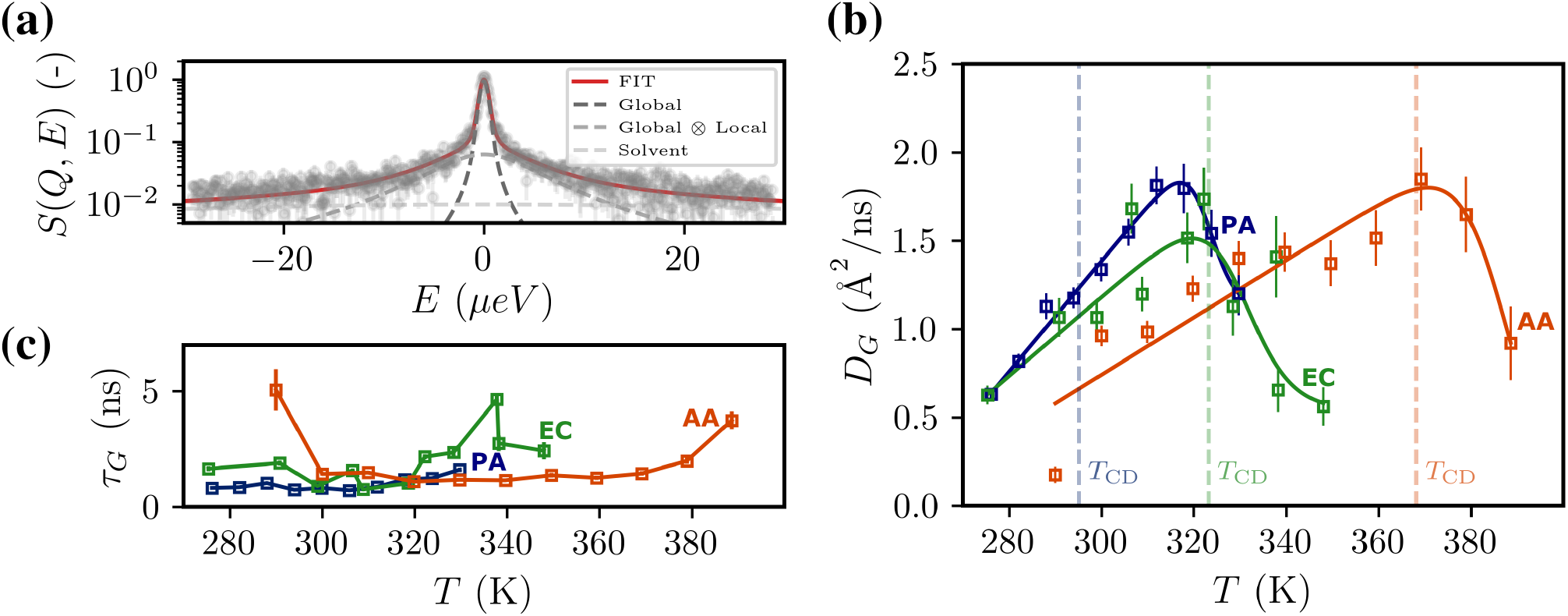
(a) Example of a fit of the scattering intensities at *Q* = 0.83 Å^−1^ and *T* = 330 K of PA. The total fit is represented by the red curve, with the Voigt components (after convolution of the Lorentzians with the resolution function) for the global and local protein contributions shown in dark gray and medium gray, respectively. The contribution of the solvent is displayed in light gray representing D_2_O. (b) Global diffusion coefficients extracted from QENS experiments of PA, EC (produced from results of [16]), and AA using the Jump Diffusion model as a function of temperature, with the respective cell death temperature *T*_CD_ shown by vertical lines. Data is fitted with the model described in Equation 2. (c) Residence times of the global diffusion according to the Jump Diffusion model for PA, EC and AA.

All spectra were fitted using the following model:

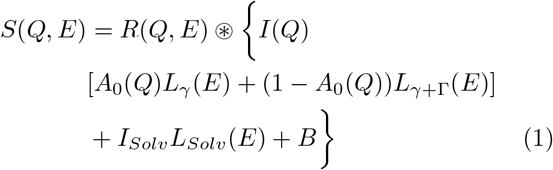

where the half width half maxima *γ* and Γ of the Lorentzian functions are characteristic for the global and local self-diffusion, respectively. The term *A*_0_ represents the elastic incoherent structure factor (EISF), *I*_*Solv*_ is the intensity of the solvent signal (in this case *D*_2_*O*), *I* is the intensity of the protein signal, *B* is the background, and *R*(*Q, E*) is the instrumental resolution function.

The global diffusion coefficients *D*_*G*_ and the time constants of diffusion *τ*_*G*_ as extracted at all temperatures using the jump-diffusion model [25] (see also Supplementary Fig. 2–4) are shown in Fig. 1b,c. These values are quite similar to the ones reported for green fluorescent protein (GFP) in EC in the range 0.3–1.4 Å^2^/ns [26, 27]. For all three bacteria the temperature dependence of the experimental *D*_*G*_ is well described by a theoretical model that captures the transition from a fluid-like to a gel-like state of the proteome [16, 28, 29]:

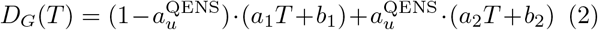

where 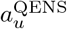 is a logistic function defined as 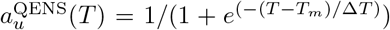. In this model, *T*_*m*_ is the midpoint of the transition, and Δ*T* describes the steepness of the curve at the transition.

A linear increase in the diffusivity as a function of the temperature can be observed for all three species at lower temperatures, in agreement with the Stokes-Einstein law [30].

However, we observe a rank among the proteomes’ diffusivity that follows the thermal stability character of the organisms. In the range 300–320 K, the diffusion coefficient *D*_*G*_ of the psychrophile is the highest, while that of the hyperthermophile is the lowest. This behavior suggests that life adaptation involves the evolutionary selection of protein diffusivity that remains constrained across the entire thermal niche. This effect is particularly evident in the hyperthermophile, which maintains a diffusivity at high temperatures comparable to that of the psychrophile and mesophile at lower ambient conditions. This supports the idea that, to maintain optimal functioning of diffusion-dependent cellular processes at elevated temperatures, overall protein diffusivity is finely regulated. These results build upon and extend previous findings showing that the root mean square atomic fluctuation amplitudes, which determine macromolecular flexibility, are similar among psychrophilic, mesophilic, and thermophilic organisms at their respective physiological [31], or death [32], temperatures.

Interestingly, protein global diffusivity reaches a maximum at an organism-specific temperature, 316.9 K for PA, 319.6 K for EC, and 370.7 K for AA. The value of the maximum diffusion coefficient falls in the range 1.5– 1.8 Å^2^/ns for all the bacteria, which suggests that there is common species-independent threshold to the diffusivity of the average bacterial protein. After such a maximum, the diffusion decreases rapidly, a behavior that we recently interpreted in the case of *E. coli* as due to the gelation of the system induced by the unfolding of a small fraction of the bacterial proteome [16]. However, while the maximum diffusivity for EC and AA occurs near their cell death temperatures *T*_CD_ i.e. 323 K and 368 K, respectively, in PA, it is notably higher than its cell death temperature of 295 K by about 22 K.

### B. Diffusion slow-down is caused by protein unfolding

In order to provide a microscopic interpretation of the dynamical arrest measured by QENS, we performed Molecular Dynamics (MD) simulations for various systems modeling the cytoplasm of the PA, and AA bacteria, and we compared the results to what already obtained for the model of the EC cytoplasm [16]. For each organism, multiple systems were set up at varying concentration levels. For each system, we created a version where all proteins were folded and one where all were unfolded. All the systems were simulated at different temperatures across the cell-death temperature of the reference organism. The PA and AA systems were assembled by considering homologous proteins of those used to model the EC cytoplasm in a previous work [16]. The detailed composition of the systems is reported in Supplementary Tab. 2–4, a molecular view is depicted in Fig. 2a.

**FIG. 2:**
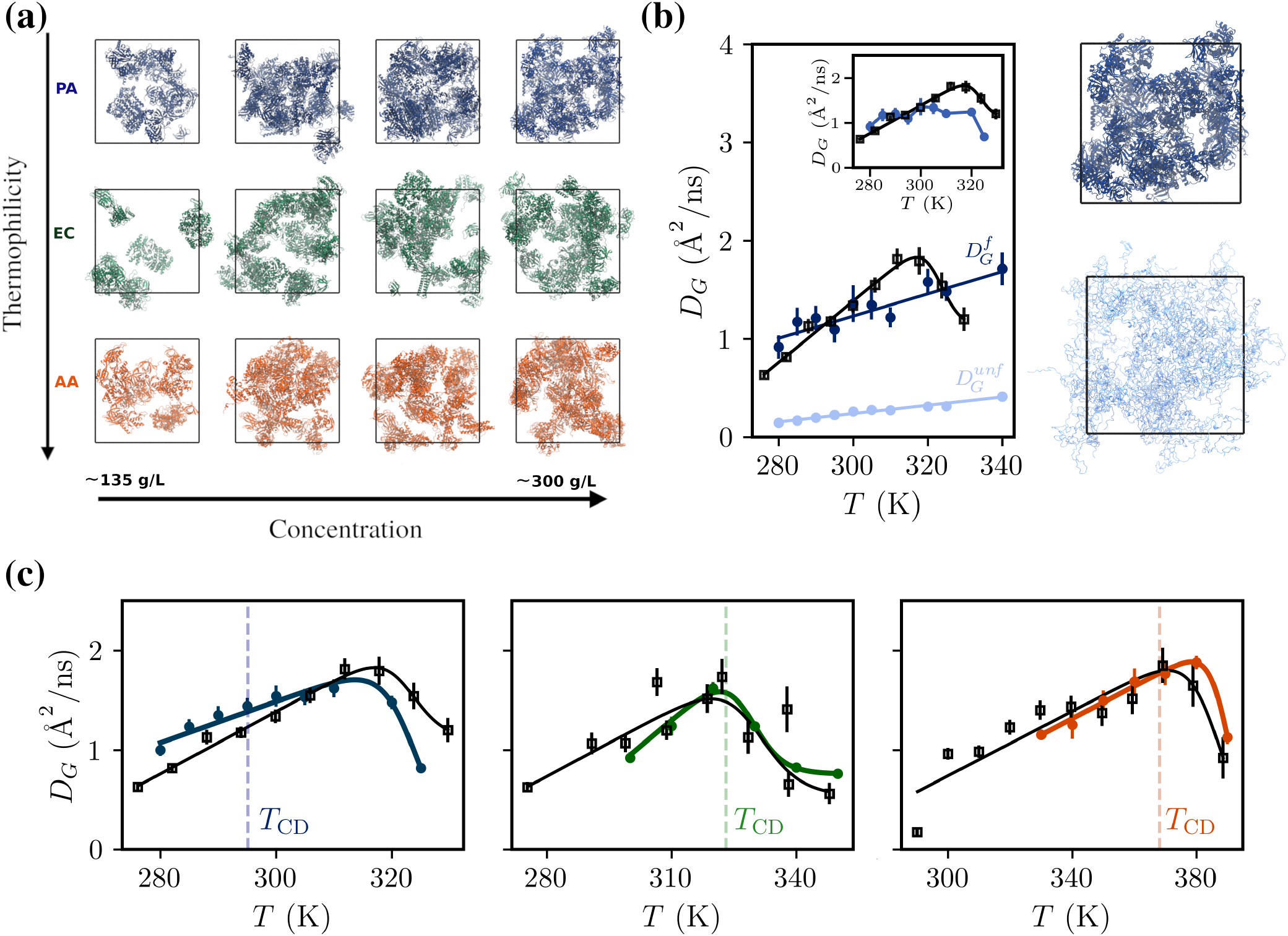
(a) Representation of the molecular systems simulated by MD of PA (blue), EC (green) and AA(red) proteins at varying concentration. (b) Global diffusion coefficients (circles) of the completely folded system, 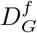 (dark blue), and the completely unfolded system, 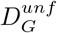 (light blue) for PA at 299 g/L which include both rotational and translational motions and fitted with a linear function. Experimental QENS data (squares) and curve is shown in black. In the inset plot, 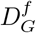 and 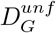 are combined (blue) according to Eq. 3 using 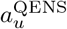 and compared with the experimental curve (black squares). (c) Optimized weighted combination of the combined diffusion of MD systems for PA (left, blue), EC (middle, green), and AA (right, orange) at different concentrations, chosen to best match the experimental QENS curves (black), allowing the deduction of the optimal concentration level.

To quantify the diffusive motions of folded and unfolded proteins, the translational and rotational diffusion coefficients of the completely folded and unfolded systems have been computed and combined to obtain a global diffusion coefficient (*D*_*G*_) to be compared directly with the experimental measurements. For sake of example, we report in Fig. 2b the temperature variation of the *D*_*G*_ computed for a completely folded 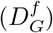 and a completely unfolded system 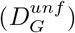 modeling the PA cytoplasm at a concentration of 299 g/L. In the figure we also report the QENS experimental values extracted from QENS measured for PA (black squares). The results for all the simulated systems, PA, EC, and AA, at different concentrations, are reported in Supplementary Fig. 6. In all the systems, as expected, we find a much lower diffusion coefficient upon unfolding of the proteins. In fact, when the proteins are unfolded, they form an entangled system characterized by a high number of protein-protein contacts [16] that contribute to increasing the system viscosity and slowing down protein mobility.

The folded and unfolded diffusion coefficients can, at each temperature, be combined together in order to reproduce the experimental data and by assuming a linear dependence on the apparent fraction of unfolded proteins 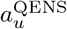 in the systems:

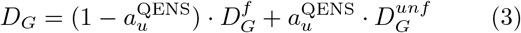

where the coefficient 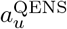 is temperature dependent and is deduced by fitting the experimental data as discussed in the previous session. For the PA system at 299 g/L the result of this combination is shown in the inset plot of Fig. 2b (see also Supplementary Fig. 7).

It should be noted that the concentration level of the simulated protein solutions considerably affects the computed value of diffusion coefficients for both the folded and unfolded systems. Therefore, the match with experimental QENS value would depend on the chosen concentration state. We have designed an optimization procedure to combine all the simulative results and deduce the concentration level that allows the diffusivity from simulation to best reproduce the experimental results. In fact, the apparent global diffusion obtained for each system at different concentrations, *D*_*G*_(*c*_*i*_), can be combined using a set of weights: 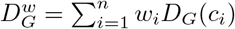. The weights can then be optimized in order to match the experimental values 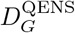:

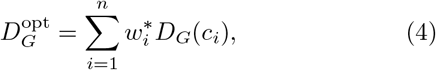

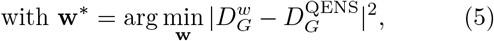

where 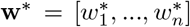 are the optimized weights and 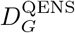 is the experimental global diffusivity that, for numerical purposes, can be interpolated at the relevant temperatures where the simulative *D*_*G*_(*c*_*i*_) are computed. The optimized curves for the three organisms are shown in Fig. 2c. They match the experimental data quite well, initially showing a thermal increase in diffusivity, followed by an abrupt slowdown. The associated optimal concentrations are 280.5 g/L, 280.0 g/L and 298.7 g/L, respectively for PA, EC and AA, showing a clear difference between the concentration of PA and EC with AA. This result is consistent with the view that the hyperthermophilic AA requires high temperatures to achieve functional protein mobility so to overcome a higher crowded and less fluid cytoplasmic state [32].

### C. Partial unfolding effect and ratio of unfolded proteins

To assess the impact of the progressive thermal protein unfolding on the diffusive dynamics, we deployed a strategy already used for EC [16]. We built, for PA and AA, three extra systems at a given concentration (299 g/L) with a varying content of unfolded proteins (25%, 50% and 75%). The proteins that have been unfolded were selected randomly (see Supplementary Tab. 5–7). From the simulations of these systems, we extracted the average translational diffusion of the proteins as a function of temperature that we report in Fig. 3a,d. By increasing the ratio of unfolded proteins, the diffusivity decreases at all simulated temperatures but not linearly. In particular, for PA already the unfolding of 25% of the proteins can cause a significant decrease in the diffusion. The slowdown caused by the progressive unfolding is related to the increase in inter-protein contacts and the shift toward a more highly viscous, entangled state. As shown in Fig. 3b,c,e,f as soon as a fraction of proteins is unfolded the probability distribution of inter-molecular contacts starts exhibiting a long power-law tail, *P* (*c*) ∝ *c*^−*k*^, and therefore a larger second moment, as indicated by the standard deviation *σ* of the distributions. The resulting entanglement corresponds to the formation of protein percolating clusters across the system, see Supplementary Fig. 10, echoing a gelation process [33]. This result extends the finding that protein diffusivity under crowded conditions is controlled by clustering dynamics [34, 35] to the case where unfolding occurs and enhances the propensity to clustering because of proteins’ larger gyration radius and extended conformations as shown in Supplementary Fig. 8. A visualization of the increase of inter-molecular contacts as a function of the ratio of unfolding is presented in Fig. 3g.

**FIG. 3:**
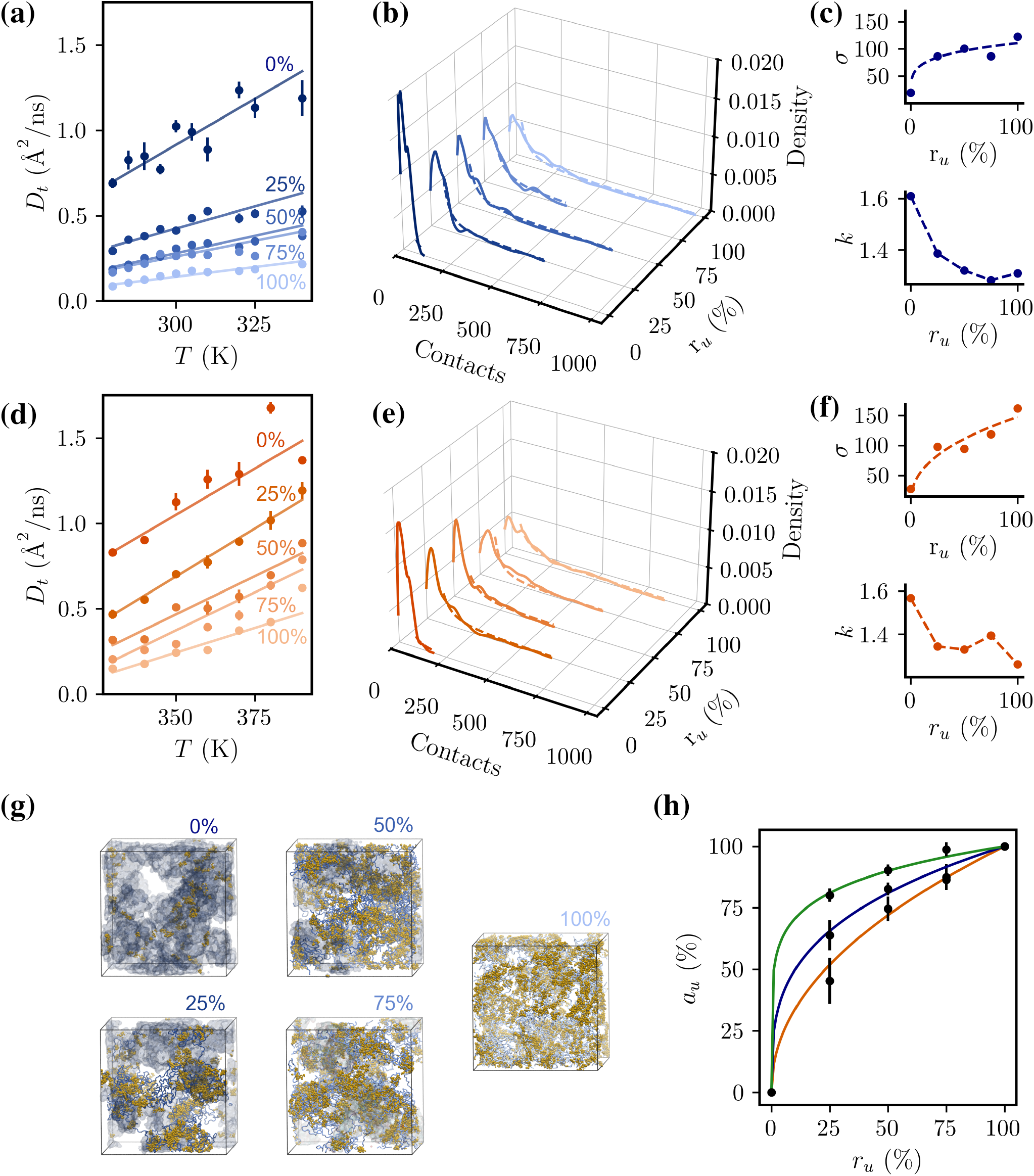
(a,d) Translational diffusion coefficient of partially unfolded systems of PA and AA at 299 g/L. (b,e) Probability distribution of inter-protein contacts as a function of the ratio of unfolded proteins for PA (at T=300 K) and AA (at T=330 K) systems. The tail of the distributions have been fitted with a power-law, *P* (*c*) ∝ *c*^−*k*^. (c,f) Standard deviation (*σ*) and power-law exponent *k* of the inter-protein contacts distributions for PA and AA system. (g) Molecular representations of simulated systems of PA (299 g/L) at different ratio of unfolding. Inter-protein contacts below 3 Å are represented in yellow. (h) Apparent fraction of unfolding *a*_*u*_ vs. ratio of unfolding *r*_*u*_ for PA (blue), EC (green) and AA (orange)

Now, following in spirit the analysis of the experimental data, at any temperature the diffusion coefficient of the partially unfolded systems can be modeled as a linear combination of the diffusion coefficients of the folded and unfolded systems, weighted by an apparent unfolded fraction coefficient *a*_*u*_. Our simulation results allow us to establish a relationship between the apparent fraction of unfolding, *a*_*u*_ (used to linearize the dependence of *D* on the unfolding progress), and the real fraction of unfolded proteins in the system, *r*_*u*_ 0.25, 0.50, 0.75. Because of the intrinsic non-linear progress seen in Fig. 3a,d, we retrieve for all systems a power law relationship 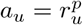, that we represent in Fig. 3h, with exponents *p* = 0.305, 0.142, and 0.474 for PA, EC, and AA, respectively. By inverting this relationship it is possible to derive from the experimental 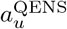, extracted from the QENS global diffusivity, the real fraction of unfolded proteins associated to the change of experimental diffusivity as a function of temperature, 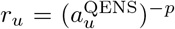. As shown in Fig. 4, for EC and AA the onset of the proteome unfolding is located a few kelvin above the *T*_CD_. These results suggest that for these mesophilic and hyperthermophilic organisms only a few proteins are actually unfolded at the cell death [16, 36, 37]. The psychrobacter PA, however, behaves surprisingly, as the proteome unfolds more than 20 K above the predicted cell-death temperature. This indicates that, for this psychrophilic organism, other molecular processes cause the thermal cell death.

**FIG. 4:**
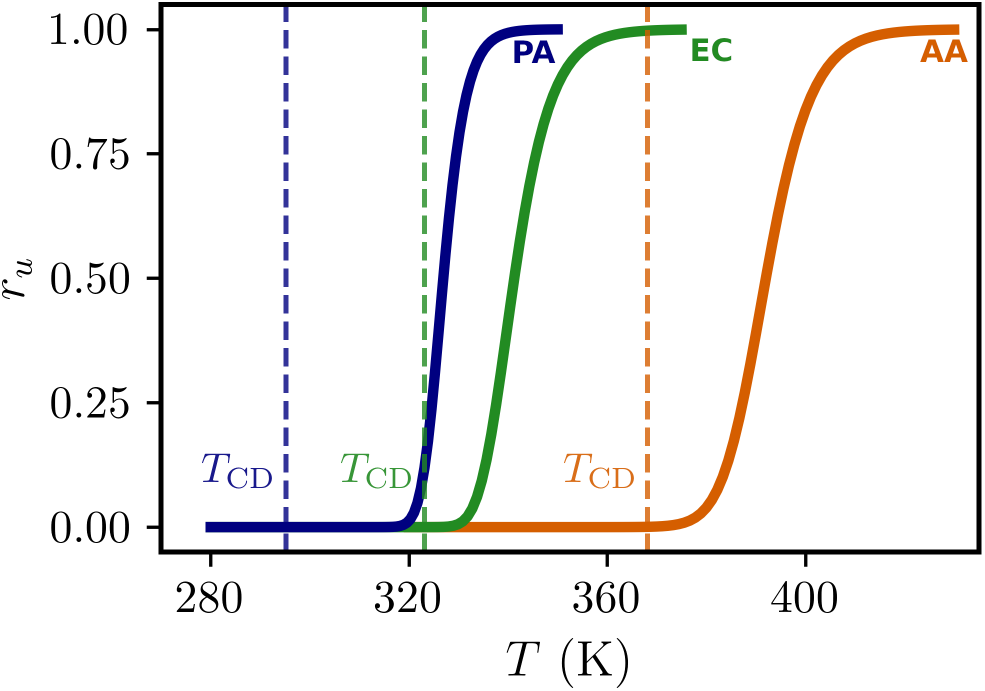
Ratio of unfolded proteins as a function of temperature from QENS experiments and MD simulations of PA (left), EC (middle), and AA (right). Vertical lines show the respective cell death temperature of the organism. The arrows indicate the ratio of unfolded proteins at that temperature. The results indicate a strong resistance to temperature of the psychrophilic proteome, which upholds 24 K beyond the cellular death.

### D. Protein melting temperature prediction and DSC measurements show the high stability of *P. arcticus* proteome

First, we want to confirm that the unfolding onset of the proteome of PA deduced by the temperature dependence of the dynamical properties of the proteome, really occurs at a temperature close to 320 K, and therefore more than 20 K beyond the cell death of the organism. Differential Scanning Calorimetry (DSC) measurements of the whole cells have been performed for PA and EC.

In Fig. 5a (top) we plot the behavior of the measured heat capacity *C*_*p*_ as a function of temperature. While for a protein in dilute solution the measured peak of heat flux is attributed to the transition from a folded to an unfolded state [38], in the case of a whole cell, the mixture of proteins, nucleic acids, and membrane lipids can undergo multiple transitions [39], each contributing to the global heat flux. The signal of DSC of a whole cell is therefore expected to reflect the sum of these contributions, resulting in a broader profile with multiple peaks.

**FIG. 5:**
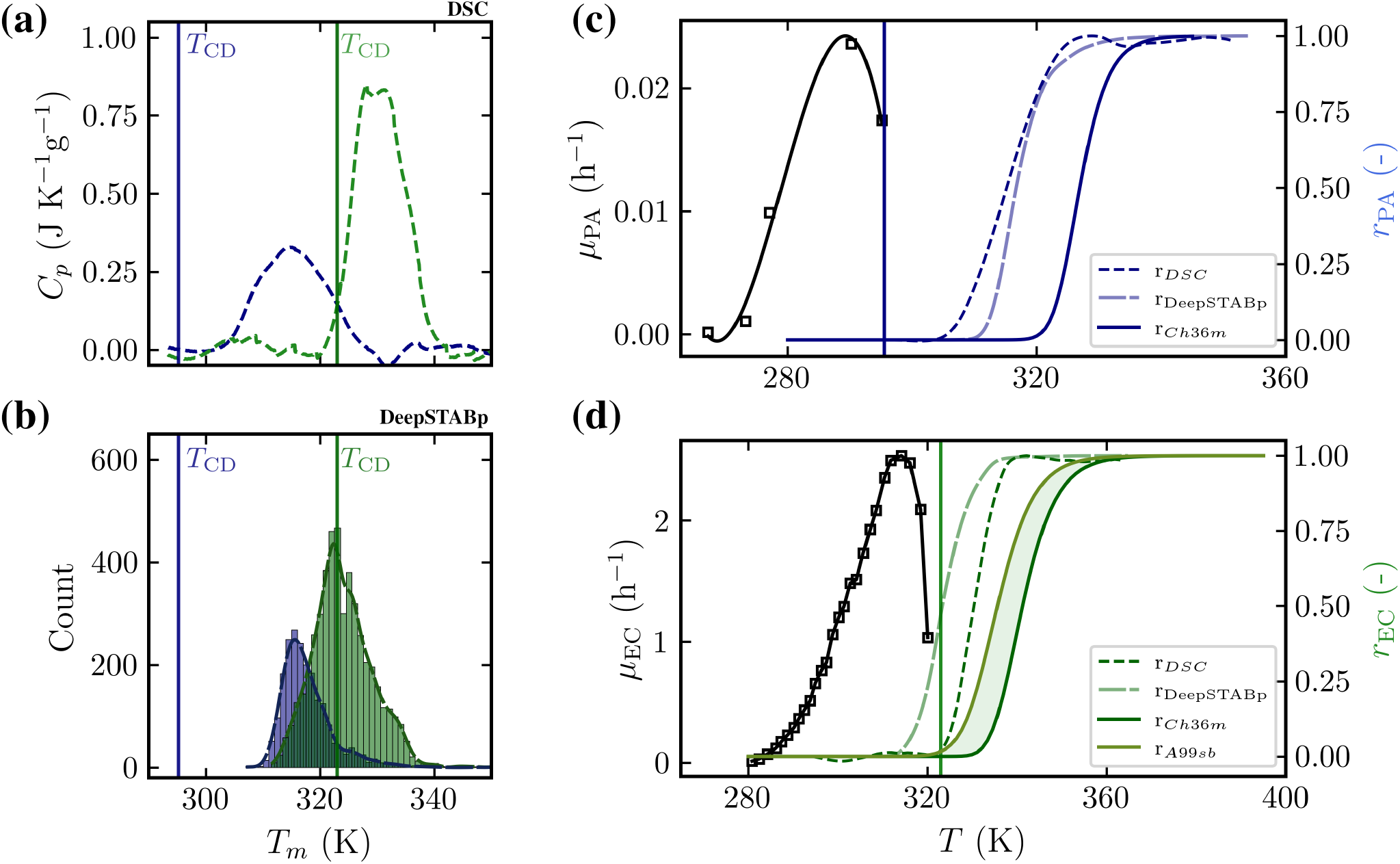
(a) Heat Capacity from DSC experiments of PA and EC with the respective cell death temperature as vertical lines for PA (blue) and EC (green). (b) Distributions of the predicted melting temperatures of PA and EC proteins sequences from UniProt using the DeepSTABp tool [45] for PA (blue) and EC (green). (c) Experimental growth rate curve (black squares) from [46] with a polynomial fit showing its trend and predicted fraction of unfolded proteins from DSC, DeepSTABp and MD+QENS methods in PA. (d) Experimental growth rate curve (black squares) from [47] and predicted fraction of unfolded proteins from DSC, DeepSTABp and MD+QENS methods using the Amber99sb-disp force field and Charmm36m force field in EC.

The curve associated to EC is in very good agreement with previous DSC scans reported in literature, [39–41] with a main peak present between 320 K and 338 K. Interestingly, the onset of this main peak corresponds to the thermal death of EC (322 K). A similar main peak is observed for the PA cells in the temperature range 303-326 K. Smaller peaks are following at higher temperatures of 336 K and 343 K, as also found and described by Mackey *et al*. [39] for other organisms. The origin of the main peak is still debated, with some attributing it solely to ribosomes and rRNA degradation [39], while others relate it more generally to the sum of individual protein unfolding events [42]. While for EC, the comparison between the temperature range of the main DSC peak and the set of melting temperatures of individual EC proteins is supported by a large body of literature [12, 36, 40, 41], for PA, this task is more difficult since experimental data are more scarce. A recent study on the DNA-binding protein ParI from PA describes a melting beginning at 321 K [43]. Similarly, Nowak *et al*. found a *T*_*m*_ of 333 K for another DNA-binding protein of PA with DSC [44]. To provide complementary support, we exploit a machine learning-based melting temperature prediction tool, DeepSTABp [45], which allows us to deduce the melting temperature of a large set of proteins from PA and EC (see Methods section IV G). This model was trained on millions of protein sequences, incorporating organism-specific features such as the presence of ions and chaperones, and differentiating between cell lysates and whole cell measurements. The predicted melting temperatures are plotted in Fig. 5b. The distribution of the melting temperatures [48] for the PA proteins has at 315.6 K a maximum in agreement with the DSC thermogram peak (314.7 K) and shows a long tail extending up to 351 K. The distribution of EC proteins’ melting temperatures is more symmetric, with a maximum at a temperature of 322.3 K, again close to the one of the main DSC peak (328.1 K). These results clearly support the interpretation proposed by Lepock *et al*. [42], thus suggesting that the broad DSC profile originates from the overlap of individual protein unfolding events. On this basis, we integrated the DSC curves to retrieve, after normalization, the fraction of unfolded proteins as a function of the temperature [42]. This fraction can be directly compared to the analogous quantity obtained by combining QENS experiments with MD simulations. A comparison with available DSC data was also proposed by Gosh and Dill when discussing their theoretical model describing the physical limit of proteome stability for different organisms [48, 49]. The results are shown in Fig. 5c,d for PA and EC. In general, we observe that the unfolding rates obtained from QENS/MD are shifted approximately 10 K higher than those derived from DSC, while maintaining a similar cooperative trend. We should also emphasize that, as discussed in our previous work [16], the choice of force field affects the fine details of the curve extracted from QENS/MD, as shown in Fig. 5d, where we compare results for the EC proteome simulated with the CHARMM36m [50] and AMBER a99SB-disp [51] force fields. Interestingly, the fractions of unfolded proteins calculated from the distribution of melting temperatures are in close agreement with those derived from DSC, especially in the case of PA. When comparing the fraction of unfolded proteins with the bacterial growth rate curve reproduced from [46, 47], we confirm that while in EC cell death is associated with the onset of proteome melting, in PA, proteome unfolding occurs at a significantly higher temperature than its cell death.

## III. DISCUSSION AND CONCLUSION

In this study, we combined QENS experiments and MD simulations to investigate the nanosecond-scale diffusion dynamics of the proteome in psychrophilic, mesophilic, and hyperthermophilic bacteria. QENS experiments show that for the three species we observe a similar temperature dependence of the diffusivity of proteins in the cytoplasm. As we have previously shown [16], translational motion contributes for more than 80% to *D*_*G*_, therefore its initial linear increase is likely related to a Stokes-Einstein-like regime of proteins diffusing in a crowded milieu. Then, after peaking at a critical temperature, the diffusivity abruptly slows down.MD simulations show that this dynamical arrest is linked to the onset of proteome unfolding in the cytoplasm. Even the unfolding of a relatively small fraction of the proteome is accompanied by strong inter-protein entanglement, which in turn increases cytoplasmic viscosity.

Quite interestingly, the global dynamics of the three proteomes exhibit a hierarchical ranking in the room temperature range, with the extent of diffusive motions progressively decreasing from psychrophilic to mesophilic to hyperthermophilic. This finding, supported by direct viscosity calculations, suggests that adaptation to cold temperatures benefits from a more fluid cytoplasmic state, compensating for reduced thermal fluctuations in low-temperature environments. On the other hand, hyperthermophiles require high thermal excitations to achieve the same level of fluidity as psychrophilic or mesophilic bacteria. From an evolutionary perspective, the relatively high cytoplasmic viscosity in hyperthermophiles has been suggested as a mechanism to reduce the thermal degradation of nicotinamide adenine dinucleotide (NADH) [52]. Additionally, by matching the diffusivity calculated from simulations with the experimental QENS results, we estimated the optimal cytoplasmic concentration for the three species, which again exhibit a hierarchical ranking, from the less crowded state of psychrophilic PA (280 g/L) to the more crowded condition of hyperthermophilic AA (298 g/L). However, this small variable concentration can not explain the marked differences in proteome mobility observed for the three organisms at a given temperature. The functional fluidity of the psychrophilic cytoplasm would mainly arise from an evolutionary adaptation of the amino acid composition of proteins [53]—enriched in negatively charged residues—which, at a given concentration, facilitates faster motion [27, 32, 54, 55]. On the other hand, a less negative charge associated with the average protein and stickier inter-protein interactions would lead to slower diffusive motions in hyperthermophilic bacteria.

EC and AA attain very similar values of the maximum attainable diffusion coefficients in proximity of their optimal growth temperatures (~ 319 K for EC and ~358 K for AA). This finding suggests that the stochastic diffusive dynamics of the proteins in the bacterial cytoplasm have been optimized together with the common micrometric cellular size [56] and the crowding conditions of biomolecules [49] to allow for the covering of the cytoplasmic volume in characteristic times of the order of 10^−2^–10^−1^ s [49]. However, the fact that, in the case of PA, cell death occurs well before the maximum diffusion coefficient is attained suggests that the optimization of global diffusive dynamics is a necessary but not sufficient condition for cellular growth, especially in cold environments. Indeed, in proximity of the cell-death temperature for EC and AA a marked slowdown of the protein global diffusion is observed, and for both organisms our data support the view that only a small set of essential proteins unfold at this temperature [36, 37]. In stark contrast, PA exhibits a markedly different behavior, with the Stokes-Einstein trend remaining largely unaffected until approximately 20 K above the cell-death temperature. DSC experiments performed on whole PA cells, along with machine-learning predictions of protein melting [45], confirm that the signature of proteome unfolding coincides with the observed dynamical arrest, highlighting a significant decoupling between cellular viability and proteome stability. We thus encounter a striking anomaly: the metabolism of the cold-adapted PA bacterium ceases long before the structural integrity of its protein machinery begins to degrade. We therefore provide evidence that the temperature range of cell viability in psychrophilic bacteria is fundamentally decoupled from the thermal stability of the proteome. This finding challenges the prevailing assumption that cellular survival is inherently tied to proteome integrity, revealing instead a striking disconnect between metabolic viability and structural stability. Notably, in the cases of the psychrophilic bacteria *Bacillus psychrophilus* and *Oleispira antarctica*, a similar shift between the thermal cell death temperature and the onset of the main calorimetric peak has been observed [42]. However, no conclusions were drawn in that study regarding the decoupling between metabolic functionality and proteome structural stability, a key aspect that we highlight here for the first time. A large body of literature has been devoted to explain why the bacterial growth-rate reaches a maximum as a function of temperature and then drastically decreases [57]. The commonly accepted view is that at high enough temperatures a non-negligible fraction of proteins unfolds, some of which are enzymes critical for the bacterial metabolism, which causes the cell-death [49, 57, 58]. Our data questions that a similar view holds for psychrophilic bacteria. In these bacteria, thermal death occurs due to a cascade of destabilizing events: proteome composition and metabolic collapse, membrane lipid disorganization, oxidative stress and nucleic acid damage [59, 60]. These organisms’ adaptations to cold – enhanced enzyme flexibility, fluid membranes, and specialized repair systems – render them exquisitely sensitive to mild warming. On these grounds, both the principle of structure–function stability trade-off and the presumed direct correlation between proteome stability and cellular viability are deeply challenged in the case of psychrophiles.

Strikingly, the observed shift between the optimal growth temperature (or the close cell-death temperature) and the temperature of proteome melting echoes what was already observed for individual enzymes from psychrophilic organisms [61–63]. For several of these enzymes, it has been shown that they lose their activity at significantly lower temperatures, well before the onset of their unfolding [62, 64]. Several models have been proposed to explain the origin of this gap between the optimal catalytic temperature, *T*_*opt*_, and the unfolding temperature *T*_*m*_ [65–67]. For instance, psychrophilic *α*-amylases lose activity at temperatures before unfolding, a process attributed to heat-induced distortions in the active site and the existence of dead-end catalytic configurations [66]. This localized dysfunction suggests that critical enzymatic processes fail before bulk protein denaturation becomes detectable. A more general view was offered by comparing the experimental catalytic efficiency as a function of temperature between two homologous dihydrofolate reductase enzymes from mesophilic and thermophilic organisms [67]. It was shown that the gap between *T*_*opt*_ and *T*_*m*_-that is different among the two homologues- can be rationalized and quantified by comparing the activation energy *E*_*a*_ of the catalytic activity of the two enzymes and their thermodynamic unfolding enthalpies, Δ*H*_*u*_ [67]. In order to link the observed decoupling between metabolism and proteome stability and diffusion to single enzyme behavior, a crucial next step will be the *in-vivo* investigation of the temperature-dependent relationship between activity and stability, as well as the local dynamics of active sites.

Moreover, in order to cope with cold psychrophilic bacteria over-express specific enzymes involved in transcription and translation [68, 69] and produce abundant cryprotecting metabolites. Our result suggests that above a critical temperature the proteome composition of psychrophilic bacteria could change toward a folded but unfunctional state unable to ensure metabolism. Precise temperature-dependent proteome profiling of cold-adapted bacteria are needed to provide a decisive answer to this open question.

In conclusion, our work provides valuable insights into the hierarchical thermal vulnerability of different bacterial families. This is key to informing strategies for food preservation, bioremediation, and the sustainable use of cold-adapted microbes in biotechnology. Future research should explore evolutionary trade-offs in psychrophiles’ heat sensitivity and engineer synthetic variants with stabilized enzymes for industrial applications.

## IV. METHODS

### A. Sample preparation for neutron scattering experiments

We investigated three bacteria with varying thermal stabilities: the psychrophilic *Psychrobacter arcticus* 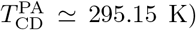 [47], the mesophilic *Escherichia coli* 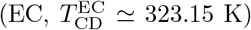 [49], and the hyperthermophilic *Aquifex aeolicus* 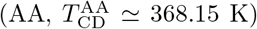 [70]. EC neutron scattering data was already available from [16].

*Aquifex aeolicus* VF5 was cultivated at 85 °C following the method previously described in [71], using SME modified medium. Cultures were prepared in 2 L bottles at pH 7.3, with sodium thiosulfate present and under a H_2_/CO_2_/O_2_ atmosphere. *Psychrobacter arcticus* cells (DSM 17307) were obtained after being grown for three days at 17 °C with shaking (150 rpm) in Bacto Marine broth (DIFCO 2216, DSM Medium 514), reaching a final OD_600nm_ of 1.5. In both cases, the cells were cultivated in media prepared with H_2_O, harvested during the late exponential growth phase by centrifugation, and washed twice with a buffer at pD 8 (50 mM Tris, 150 mM NaCl, 5 mM KCl) prepared with D_2_O (99.9 atom D), as described in [16]. To achieve a pD of 8, the buffer pH was adjusted to 7.6 using HCl. Finally, 1.5 g and 2.3 g of washed, unfrozen cells of *A. aeolicus* and *P. arcticus* were obtained and analyzed.

### B. Quasi-elastic incoherent neutron scattering experiments

QENS experiments were carried out using the IN16b spectrometer at the Institut Laue-Langevin (ILL) in Grenoble [23] (DOI: 10.5291/ILL-DATA.8-04-960). Data were collected with an energy resolution of 0.9 *μ*eV, corresponding to a time window ranging from 0.3 to 5 ns. The experiments were conducted on samples containing whole-cell organisms, as well as on a vanadium sample and an empty cell. Measurements were performed over various temperature ranges to encompass both the organisms’ optimal growth conditions and their upper thermal stability limits: from 276 K to 324 K in 6 K increments for PA and from 290 K to 390 K in 10 K steps for AA.

### C. Neutron Data Analysis

QENS data were analyzed using a common model for protein diffusion from [22, 29].

Generally, the experimental scattering signal is obtained by convolving the instrumental resolution function, *R*(*Q, E*), which is determined through vanadium measurements, with a theoretical model. Here *R*(*Q, E*) was fitted by a sum of a Gaussian function and a constant background as shown in Supplementary Fig. 1. Since proteins have the highest proportion of hydrogen atoms (and thus have a high scattering cross-section) and constitute more than half of the dry weight in bacterial cells, they are expected to dominate the theoretical function [15]. As a result, the theoretical function is commonly modelled by

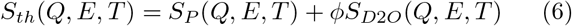

where *S*_*P*_ (*Q, E, T*) corresponds to the scattering contribution from cellular proteins, and 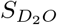 (*Q, E, T*) represents the contribution from the *D*_2_*O* buffer surrounding the cells. The parameter *ϕ* denotes the fraction of *D*_2_*O* in the samples. The *D*_2_*O* signal, 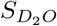, can be accurately described using a Lorentzian function [29]. To simplify the model, parameters for the Lorentzian function were estimated based on prior measurements of *D*_2_*O* and *D*_2_*O* + PBS samples, see [16]. The final model of the scattering function *S*(*Q, E*) therefore takes the analytical form presented in Equation 1. The fitting process was conducted in two sequential steps, involving a normal fit and a simultaneous fit, as further described in Supplementary Note 1 and Supplementary Fig. 3.

### D. MD systems

All-atom MD simulations were conducted in this study to investigate the dynamics of cytoplasm models of the three bacteria, PA, EC, and AA.

Starting from a set of systems used to investigate the dynamics of EC cytoplasm in a previous work [16], we built here equivalent systems for the PA and AA organisms by considering homologous proteins and similar concentrations (see Supplementary Tab. 2-4).

Where available, the 3D pdb structure from the PDB databank was used. If the experimental structure was not accessible, an AlphaFold structure was generated and used [72]. For all proteins, for multi-domain proteins, we assumed the same oligomeric state as the corresponding EC protein. Before assembling the proteins into the cytoplasm model, their structures were relaxed via an equilibration procedure using GROMACS [73], as explained in Supplementary Note 2. The Packmol software [74, 75] was used to place the equilibrated proteins into a 170 Å cubic box with some final manual corrections to avoid clashes. The number of inserted proteins depends on the target concentration.

### E. MD simulations

All simulations were performed using GROMACS versions 2019.4 and 2021.3 [73], employing the Charmm36m force field [50]. Water was modeled using the TIP3P model [76]. The boxes were subsequently hydrated, and 150 mM of K^+^ and Cl^−^ ions were added, including some extra ions to neutralize the net charge of the system. The systems underwent energy minimization, reducing the force constant to below 1000 kJ mol^−1^ nm^−1^. For all the systems containing the folded proteins, a 6-step NVT procedure lasting 11 ns using the V-rescale thermostat [77] with decreasing position restraints was applied to the proteins in the system, and a 100 ns NPT equilibration with the Berendsen barostat [78] was subsequently run. In parallel, with the equilibration of the systems of folded proteins, a protocol of unfolding was devised to create the same systems with completely unfolded proteins. The unfolding process involved exposing the proteins to elevated temperatures (1200 K for PA and 1500 K for EC and AA), see Supplementary Fig. 5. After this high-temperature treatment, a cooling protocol was applied to bring the systems back to the same temperatures as the folded counterparts. These unfolded systems were then equilibrated using the same procedure as for the folded proteins. For each organism, partial unfolding was performed on one system. Three additional states of partial unfolding were also created, corresponding to unfolding fractions *r*_*u*_ = 25, 50, 75 %, see Supplementary Tab. 5-7. This was achieved by applying the same protocol used for the fully unfolded systems, but with the respective percentages of folded proteins remaining frozen during the unfolding and cooling steps (see also [16]). Proteins selected for unfolding were chosen randomly to match the desired unfolding rates, which were determined based on the fraction of hydrogen atoms in each protein.

A 102.4 ns production run was performed in the NPT ensemble using the Parrinello-Rahman barostat [79], with temperatures set according to the relevant ranges for each organism. For the steps outlined above, a cutoff of 1.2 nm was applied for short-range non-bonded interactions. Van der Waals forces were smoothly reduced to zero between 1.0 and 1.2 nm. Long-range electrostatic interactions were calculated using the particle mesh Ewald (PME) method [80]. The Leap-Frog algorithm was used to propagate Newton’s equations of motion. The LINCS algorithm was applied to constrain the lengths of all protein bonds involving hydrogen [81], while the SETTLE algorithm was employed to maintain the rigidity of water molecules [82].

### F. MD Data Analysis

A comparable quantity as the experimental apparent global diffusion can be extracted from the MD data. The translational and rotational diffusion coefficients were evaluated for all the protein chains at the relevant time window of 0.3 to 5 ns corresponding to the resolution of the experimental measurements. The translational diffusion was calculated by following the protein’s center of mass and by calculating the mean-square-displacement (MSD). The MSD was linearly fitted in the 0.3-5 ns time region. For the rotational diffusion, 1000 random unit vectors were generated in the protein structure. At each timestep, the protein chains were centered and rotated with respect to a reference initial structure to estimate a rotation matrix. The rotation of each vector was then estimated by applying the rotation matrix. The rotational autocorrelation function was then calculated using a second-order Legendre polynomial. By fitting the rotational autocorrelation with an exponential decay function in the same time regime as the translational diffusion, from 0.3 to 5 ns, a decay time *τ*_*r*_ was obtained. The rotational diffusion coefficient was derived as *D*_*r*_ = 1/(6*τ*_*r*_), for each chain in all the systems and all temperatures. The rotational and translational coefficients were finally corrected for PBC conditions [83, 84] using the viscosity calculated as explained in Supplementary Note 3. To obtain the global diffusion coefficient for comparison with the experiments we used the relation

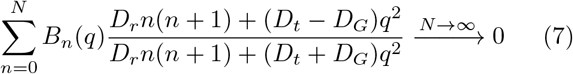

with

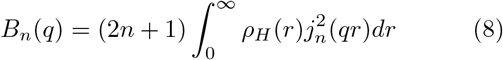

where *ρ*_*H*_ is the radial distribution function of the chain’s hydrogen atoms and *j*_*n*_ is the spherical Bessel function of order n. To this end, *N* was fixed to a sufficiently large number to ensure convergence with *N* = 550. Equation 7 was solved numerically using a minimization problem, where we set *q* to *q* = 2. To compare the results with the experimentally extracted global diffusion, *D*_*G*_ was weighted by the hydrogen atoms of each chain. In this way, for each system at each temperature, one single *D*_*G*_ value was obtained.

Further details on protein radii of gyration and proteinprotein inter-molecular contacts are provided in Supplementary Fig. 8,9.

### G. Melting Temperature Prediction

To predict the melting temperatures of various proteins from PA, EC, AA, the open-source tool DeepSTABp was used [45]. The growth temperatures used were the same as those for the preparation of the experimental samples of the respective bacteria: 17 °C for PA, 37 °C for EC, and 85 °C for AA. For the predictions, the cellular environment was selected. All proteins available in the UniProt database for the respective bacteria were included, resulting in a total of 4850 proteins for PA, 2103 proteins for EC, and 5308 proteins for AA.

### H. DSC measurements

Differential Scanning Calorimetry (DSC) measurements of the whole cells of PA and EC with the same samples as used in the QENS experiments were performed with a multicell instrument from TA at ILL, Grenoble. The experiment consisted of one reference cell filled with the buffer as in QENS, while the other cells were filled with bacterial samples and buffer. The machine was heated up from 20 °C to 90 °C (the maximum possible) and then heated down again to 20 °C. The DSC measurements were all analysed using the open-source code pyDSC [85]. The weights of the samples were 538.8, 512.3 and 537.1 g with a concentration of 26, 20 and 24 mg/ml for PA, EC, and AA, respectively.

## ACKNOWLEDGMENTS

F.S. and B.C. acknowledge the financial support by the “Initiative d’Excellence” program from the French State (Grant “DYNAMO”, ANR-11-LABX-0011-01, and “CACSICE”, ANR-11-EQPX-0008). Part of this work was performed using HPC resources from GENCI [CINES, TGCC, IDRIS] (grant x2023(4)6818) and LBT. S.T. acknowledges funding support by the Czech Academy of Sciences (Lumina Quaeruntur Fellowship LQ200402301). The authors thank the ILL for the attribution of beam time.

## AUTHOR CONTRIBUTIONS

This study is part of the Ph.D. project of B.C., carried out under the joint supervision of J.P., A.P. and F.S. J.P., F.S. and A.P. designed research; M.G. and M.T.G.O. contributed samples; J.P. T.S. and C.B. carried out experiments, with B.C. contributing. B.C. and S.T. performed simulations; B.C. analyzed data; F.S., J.P. and A.P. jointly supervised the data analysis. B.C., J.P., F.S. and A.P. wrote the paper.

## SUPPLEMENTAL INFORMATION

It contains additional details on neutron scattering experiments, molecular dynamics simulations, data analysis, and supplementary figures and tables.

