## supplemental information for "Cytoplasmic Fluidity and the Cold Life: Proteome Stability is Decoupled from Viability in Psychrophiles"

Corresponding author(s): *Double-anonymous peer review submissions: write DAPR and your manuscript number here instead of author names.*

Last updated by author(s): *YYYY-MM-DD*

### Reporting Summary

Nature Portfolio wishes to improve the reproducibility of the work that we publish. This form provides structure for consistency and transparency in reporting. For further information on Nature Portfolio policies, see our Editorial Policies and the Editorial Policy Checklist.

#### Statistics

For all statistical analyses, confirm that the following items are present in the figure legend, table legend, main text, or Methods section.

| n/a | Confirmed | |
|---|---|---|
| ☐ | ☒ | The exact sample size (*n*) for each experimental group/condition, given as a discrete number and unit of measurement |
| ☒ | ☐ | A statement on whether measurements were taken from distinct samples or whether the same sample was measured repeatedly |
| ☒ | ☐ | The statistical test(s) used AND whether they are one- or two-sided<br>*Only common tests should be described solely by name; describe more complex techniques in the Methods section.* |
| ☒ | ☐ | A description of all covariates tested |
| ☒ | ☐ | A description of any assumptions or corrections, such as tests of normality and adjustment for multiple comparisons |
| ☒ | ☐ | A full description of the statistical parameters including central tendency (e.g. means) or other basic estimates (e.g. regression coefficient) AND variation (e.g. standard deviation) or associated estimates of uncertainty (e.g. confidence intervals) |
| ☒ | ☐ | For null hypothesis testing, the test statistic (e.g. *F*, *t*, *r*) with confidence intervals, effect sizes, degrees of freedom and *P* value noted<br>*Give P values as exact values whenever suitable.* |
| ☒ | ☐ | For Bayesian analysis, information on the choice of priors and Markov chain Monte Carlo settings |
| ☒ | ☐ | For hierarchical and complex designs, identification of the appropriate level for tests and full reporting of outcomes |
| ☒ | ☐ | Estimates of effect sizes (e.g. Cohen's *d*, Pearson's *r*), indicating how they were calculated |

*Our web collection on statistics for biologists contains articles on many of the points above.*

#### Software and code

Policy information about availability of computer code

| | |
|---|---|
| Data collection | Data were collected at ILL using the NOMAD XOP (v1.02) code (https://www.ill.eu/for-ill-users/support-labs-infrastructure/sample-environment/software/xop-plugins-for-igor-pro/nomad-xop), developed at ILL and free. |
| Data analysis | The Neutron Scattering data were reduced using Mantid, a code which was developed by ILL and which is constantly updated, and analyzed with in-house python developped code. The code will be made available. The MD simulations have been carried out using the popular gromacs MD engine. MD analysis were performed with both gromacs analysis tools or customized python scripts. |

For manuscripts utilizing custom algorithms or software that are central to the research but not yet described in published literature, software must be made available to editors and reviewers. We strongly encourage code deposition in a community repository (e.g. GitHub). See the Nature Portfolio guidelines for submitting code & software for further information.

#### Data

Policy information about availability of data

All manuscripts must include a data availability statement. This statement should provide the following information, where applicable:

- Accession codes, unique identifiers, or web links for publicly available datasets
- A description of any restrictions on data availability
- For clinical datasets or third party data, please ensure that the statement adheres to our policy

data are available after a period of embargo at the webpage data.ill.fr

#### Research involving human participants, their data, or biological material

Policy information about studies with human participants or human data. See also policy information about sex, gender (identity/presentation), and sexual orientation and race, ethnicity and racism.

| | |
|---|---|
| Reporting on sex and gender | NA |
| Reporting on race, ethnicity, or other socially relevant groupings | NA |
| Population characteristics | NA |
| Recruitment | NA |
| Ethics oversight | NA |

Note that full information on the approval of the study protocol must also be provided in the manuscript.

### Field-specific reporting

Please select the one below that is the best fit for your research. If you are not sure, read the appropriate sections before making your selection.

[x] Life sciences [ ] Behavioural & social sciences [ ] Ecological, evolutionary & environmental sciences

For a reference copy of the document with all sections, see nature.com/documents/nr-reporting-summary-flat.pdf

### Life sciences study design

All studies must disclose on these points even when the disclosure is negative.

| | |
|---|---|
| Sample size | 300 - 400 mg for each sample. |
| Data exclusions | NA |
| Replication | The experiment on E. coli was conducted two times and gave reproducible results. The experiment on A. aeolicus and P. arcticus was conducted only once. For beam time restrictions, it is almost impossible to get beam time twice for the same type of study by neutron scattering. However, neutron scattering is based on Poisson statistics and contains inherently a high repetition rate. |
| Randomization | NA |
| Blinding | NA |

### Reporting for specific materials, systems and methods

We require information from authors about some types of materials, experimental systems and methods used in many studies. Here, indicate whether each material, system or method listed is relevant to your study. If you are not sure if a list item applies to your research, read the appropriate section before selecting a response.

#### Materials & experimental systems

| n/a | Involved in the study | |
|---|---|---|
| [x] | [ ] | Antibodies |
| [x] | [ ] | Eukaryotic cell lines |
| [x] | [ ] | Palaeontology and archaeology |
| [x] | [ ] | Animals and other organisms |
| [x] | [ ] | Clinical data |
| [x] | [ ] | Dual use research of concern |
| [x] | [ ] | Plants |

#### Methods

| n/a | Involved in the study | |
|---|---|---|
| [x] | [ ] | ChIP-seq |
| [x] | [ ] | Flow cytometry |
| [x] | [ ] | MRI-based neuroimaging |

#### Plants

| | |
|---|---|
| Seed stocks | NA |
| Novel plant genotypes | NA |
| Authentication | NA |
